# *agtools*: a software framework to manipulate assembly graphs

**DOI:** 10.1101/2025.09.14.676178

**Authors:** Vijini Mallawaarachchi, George Bouras, Ryan R. Wick, Susanna R. Grigson, Bhavya Papudeshi, Robert A. Edwards

## Abstract

**Summary:** Assembly graphs are a fundamental data structure used by genome and metagenome assemblers to represent sequences and their overlap information, facilitating the assembler to construct longer genomic fragments. Apart from their core use in assemblers, assembly graphs have become increasingly important in a range of downstream applications such as metagenomic binning, plasmid detection, viral genome resolution and haplotype phasing. However, there is a need for a comprehensive tool that allows programmatic access to manipulate assembly graphs across different assembly graph formats. Here we present *agtools*, an open-source Python framework that can analyse and manipulate assembly graphs. *agtools* provides a command-line interface for tasks such as graph format conversion, segment filtering, and component extraction. It also exposes a Python package interface to load, query and analyse assembly graphs from popular genome and metagenome assemblers. This enables streamlined assembly graph-based analysis that can be integrated into other bioinformatics software and workflows.

**Availability and implementation:** The source code of *agtools* is hosted on GitHub at https://github.com/Vini2/agtools and the documentation is available at https://agtools.readthedocs.io/. *agtools* is also installable from Bioconda (https://anaconda.org/bioconda/agtools) and PyPI (https://pypi.org/project/agtools/).

## 1. Introduction

Assembly is the process of reconstructing genomes from sequencing reads and is a crucial step in many genomic and metagenomic analysis workflows. Modern assemblers typically use graph structures, collectively referred to as *assembly graphs*, to represent sequences and their connections. Assembly graphs are generated as intermediate outputs from assemblers in various formats. For example, SPAdes (Bankevich et al. 2012), Flye (Kolmogorov et al. 2019), hifiasm-meta (Feng et al. 2022) and myloasm (Shaw, Marin, and Li 2025) use the Graphical Fragment Assembly (GFA) format, MEGAHIT (Li et al. 2015) uses the FASTG format, and the String Graph Assembler (SGA) (Simpson and Durbin 2012) uses the ASQG format. Despite the differences in format, vertices (or nodes) represent sequences (or segments) and edges represent connections (or links) between the sequences (Nurk et al. 2017). Formats such as GFA represent more complex information, including *paths* and *walks* consisting of multiple sequences (https://gfa-spec.github.io/GFA-spec/GFA1.html). Terminology also varies somewhat across assemblers: non-branching paths in the assembly graph are often referred to as *unitigs* (Kececioglu and Myers 1995), and longer, optimised paths are resolved into contiguous sequences known as *contigs* (Bankevich et al. 2012). Contigs can be further combined into *scaffolds*, which are longer sequences that potentially include N bases to fill gaps (Silva et al. 2013; Wick et al. 2017). Importantly, terminology and graph definitions are not universally fixed. Different assemblers may adopt slightly different conventions for constructing graphs and representing sequences within them.

Beyond their role in genome and metagenome assembly, assembly graphs are used in many graph-based approaches for downstream applications. In metagenomic binning, assembly graphs are used to group contigs into bins that belong to different taxonomic groups based on their local connectivity (Mallawaarachchi and Lin 2022a, 2022b; Xue et al. 2022, Xue et al. 2024; Feng and Li 2024; Lamurias et al. 2022; Mallawaarachchi et al. 2024). In population genomics, assembly graphs are used for haplotype phasing as they represent alternative paths corresponding to different allelic sequences (Garg et al. 2018; Henglin et al. 2024; Kazantseva et al. 2024). In viromics, assembly graphs are used to resolve strains or variants of eukaryotic viruses and bacteriophages (Luo and Lin 2023; Mallawaarachchi et al. 2023; Papudeshi et al. 2025). Moreover, plasmids can be detected from assembly graphs as circular isolated sequences or distinct subgraphs (Rozov et al. 2017; Wickramarachchi and Lin 2022; Bouras et al. 2023; Paganini et al. 2024). In metagenomic sequence classification, assembly graphs are used to improve classification results based on neighbourhood information (Pu and Shamir 2022, 2024). These diverse applications depict the significance of assembly graphs, not just as an intermediate output from assembly, but as a valuable source of biological data and sequence connectivity.

Despite the growing interest in using assembly graphs for downstream applications, there is a limited number of tools that can capture assembler-specific graph structures while allowing users to manipulate them. Many existing tools can handle only one format and do not allow conversion between different formats. For example, tools such as gfatools (https://github.com/lh3/gfatools), PyGFA (https://github.com/pmelsted/pyGFA), RGFA (Gonnella and Kurtz 2016), GFA for Ruby (https://github.com/lmrodriguezr/gfa), GFAKluge (Dawson and Durbin 2019) and GfaPy (Gonnella and Kurtz 2017) can only parse and manipulate files in the GFA format, whereas Pyfastg (https://github.com/fedarko/pyfastg) can parse only files in the FASTG format (Supplementary Table S1). Assemblers such as ABySS (Simpson et al. 2009) include format conversion tools, but these are often assembler-specific and require installing the full software. Moreover, there is no programmatic interface that can represent and manipulate different assembly graph structures (e.g., unitig graphs and contig graphs) produced from popular assemblers such as SPAdes and Flye (Supplementary Table S1). This creates challenges for researchers who want to incorporate assembly graphs into their workflows, often requiring them to write custom code to perform graph-based analyses. Hence, there is a clear need for a software framework that allows seamless representation and manipulation of assembly graphs.

In this article, we introduce *agtools*, an open-source Python framework designed to analyse and manipulate assembly graphs. *agtools* provides a command-line interface with functionality such as filtering sequences, extracting connected components and graph format conversion. *agtools* also provides a Python package interface to represent different assembly graphs from popular assemblers. This facilitates the integration of assembly graph-based operations into larger bioinformatic workflows in a modular and reproducible manner.

## 2. Implementation and features

### 2.1 Implementation

*agtools* is implemented in the Python programming language and runs on the Linux and macOS operating systems. It provides a command-line interface with various subcommands for querying and manipulating assembly graphs, and an application programming interface (API) to represent assembly graphs for downstream analyses. The unitig- and contig-level assembly graphs are represented using python-igraph (Csardi and Nepusz 2006) and sequences are represented using Biopython (Cock et al. 2009).

*agtools* supports only the GFA 1 format (https://gfa-spec.github.io/GFA-spec/GFA1.html), which is used by many assemblers. Although a GFA 2 specification (https://gfa-spec.github.io/GFA-spec/GFA2.html) has been proposed, it has not been widely adopted by assemblers.

### 2.2 Command line interface

*agtools* provides a command-line interface for users to analyse and manipulate assembly graphs. It includes subcommands for obtaining basic information, graph manipulation and format conversion, as described in the online documentation (https://agtools.readthedocs.io/). The command line interface was developed using the Click package (https://click.palletsprojects.com/en/stable/).

### 2.3 Python package interface

*agtools* provides a Python package interface for users who want to represent and explore assembly graphs in their own bioinformatic tools and workflows. It provides an API to load graphs from GFA files. This is provided through the *UnitigGraph* class, which stores the graph as a python-igraph (Csardi and Nepusz 2006) *Graph* object, allowing *agtools* to use igraph’s built-in graph functionality for various tasks such as graph traversal, identifying connected components and obtaining neighbours of a given vertex. Unitig sequences (or segments) are stored using Biopython (Cock et al. 2009) *Seq* objects to easily obtain attributes such as length and reverse complement.

*agtools* allows users to load different assembly graphs from GFA files produced from four popular assemblers: SPAdes (Bankevich et al. 2012), MEGAHIT (Li et al. 2015) Flye (Kolmogorov et al. 2019) and myloasm (Shaw, Marin, and Li 2025). Since SPAdes and Flye provide unitig-level assembly graphs and contigs resolved from the unitig graph, *agtools* provides two functions *get_unitig_graph* and *get_contig_graph* to load the unitig graph and contig graph, respectively. MEGAHIT and myloasm produce contig-level assembly graphs with subtle differences in sequence representation which have been included in their respective *get_contig_graph* functions. If there are no assembler-specific intricacies and contig sequences are represented in the GFA files, assembly graphs from other assemblers can be loaded using the *UnitigGraph* class. Figures 1A and 1B show simple examples of loading assembly graphs using the *agtools* API. Further information about the API can be found in the online documentation (https://agtools.readthedocs.io/).

**Fig. 1.**
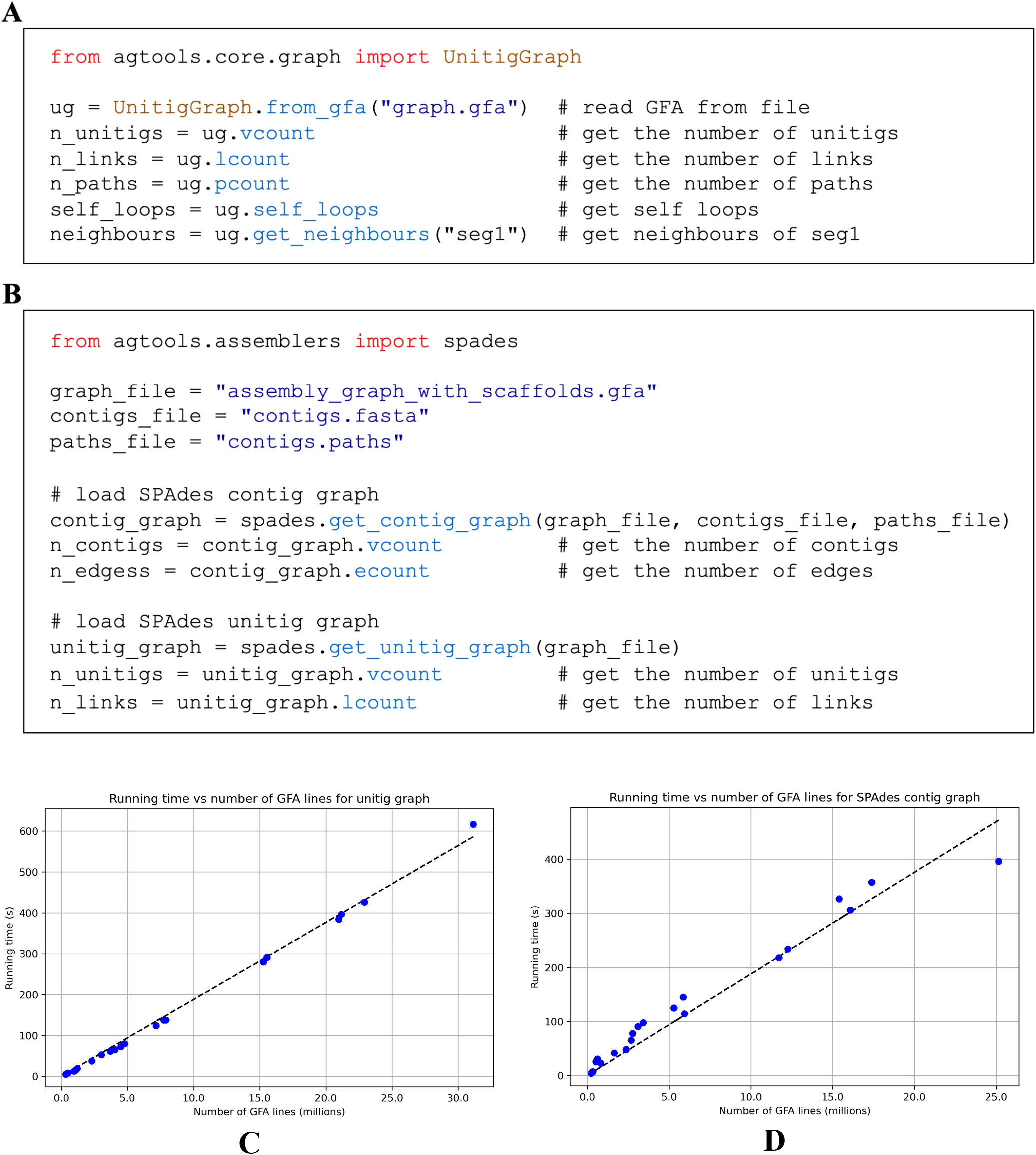
Examples of code using *agtools* to load A) unitig-level assembly graph and B) contig-level assembly graph from SPAdes assembler. Running time to load C) unitig-level assembly graphs and D) contig-level assembly graphs from SPAdes assembler vs. the number of S (segments), L (links), and P (paths) lines in the GFA file (in millions).

### 2.4 Performance enhancements

*agtools* incorporates a number of performance optimisations. When storing sequences within the *UnitigGraph* and *ContigGraph* objects, it uses an integer index instead of the original sequence identifier when building the graph which facilitates faster access. Moreover, *agtools* stores file pointers to the lines that have the sequences instead of loading the full sequences into memory. Separate functions are implemented to retrieve sequences from the GFA file in the *UnitigGraph* class and from the FASTA file in the *ContigGraph* class when required. This approach significantly reduces memory usage during graph processing.

When concatenating multiple GFA files using the *concat* subcommand, *agtools* writes the different types of lines into separate temporary files and combines them at the end, rather than keeping all the lines in memory. This approach is more memory efficient, especially when concatenating very large graph files with sizes on the order of gigabytes (refer to performance results of *concat* in Supplementary Table S4).

### 2.5 Performance analysis

We used short-read datasets from Tara Oceans (de Vargas et al. 2015) and long-read datasets from human gut metagenomes (Chen et al. 2022) and the Microflora Danica sequencing project (Sereika et al. 2025) to analyse the performance of *agtools*. The runs used from each study can be found in Supplementary Table S2. We obtained assemblies from the following assembler versions: SPAdes v3.15.5 with `--meta` flag (Nurk et al. 2017), MEGAHIT v1.2.9, Flye v2.9.6-b1802 with `--meta` flag (Kolmogorov et al. 2020) and myloasm v0.1.0. All assemblers were run with default parameters.

The unitig graph and contig graph representations of the *agtools* API were profiled using cProfile (https://docs.python.org/3/library/profile.html). Each dataset was run 10 times and the minimum, maximum, mean and standard deviation of the running time (the wall clock time between start and finish) were recorded. The CLI commands were profiled using the /usr/bin/time command. Each command was run once on each dataset. Elapsed (wall clock) time and peak memory usage (maximum resident set size) were recorded. All profiling jobs were run on a single AMD EPYC 7742 @2.25Ghz CPU node provided by the Flinders University’s DeepThought High-Performance Computing platform (Flinders University 2021). All the performance analysis results can be found in the Supplementary Material.

As evident in Fig. 1C and 1D, loading unitig graphs and contig graphs (e.g., from metaSPAdes) scales linearly with respect to the number of S (segments), L (links), and P (paths) lines in the GFA file. This indicates that the computational cost for graph loading grows in proportion to the size of the input graph representation. The performance results for loading contig graphs from the remaining assemblers MEGAHIT, Flye and myloasm can be found in the Supplementary Figures S1-S3.

### 2.6 Example applications

#### 2.6.1 Metagenomic binning

Contigs connected together in the assembly graph are more likely to belong to the same taxonomic group (Mallawaarachchi, Wickramarachchi, and Lin 2020). *agtools* can be used to group contigs based on connected components in the contig-level assembly graph into candidate bins that represent different taxonomic groups. These candidate bins can be further refined in later steps by either splitting when multiple species are in one component or merging when multiple components belong to the same species. A simple example of binning contigs based on connected components is provided in the online documentation (https://agtools.readthedocs.io/).

#### 2.6.2 Identifying plasmid candidates

Plasmids often assemble as circular segments in the assembly (Bouras et al. 2023). Hence, identifying circular within the typical range of plasmid genome length is a good starting point before proceeding to deeper validation to identify plasmids. *agtools* can be used to identify circular segments (from self-loops) in the unitig-level assembly graph. A simple example of identifying plasmid candidates is provided in the online documentation (https://agtools.readthedocs.io/).

#### 2.6.3 Identifying bacteriophage candidates

Bacteriophages with circular genomes can form circular components in the assembly graph (Mallawaarachchi et al. 2023). Hence, we can identify bacteriophage candidates by determining components that have circular genomic paths. The paths can be filtered based on the typical range of bacteriophage genome length before further validation such as scanning for highly conserved phage genes. A minimal example of identifying bacteriophage candidates is provided in the online documentation (https://agtools.readthedocs.io/).

#### 2.6.4 Haplotype phasing

Haplotype phasing is the process of identifying and reconstructing genetic variants (alleles) of haplotypes in diploid or polyploid genomes. This can be achieved by resolving *bubbles* in assembly graphs where a vertex splits into multiple outgoing edges that later reconverge into another vertex (Weisenfeld et al. 2017; Garg et al. 2018). These alternative paths in the graph correspond to sequence differences between haplotypes. A naïve example of haplotype phasing using the assembly graph is provided in the online documentation (https://agtools.readthedocs.io/).

## 3. Conclusion

*agtools* is an open-source Python framework that can analyse and manipulate assembly graphs. It provides a command-line interface to query and manipulate assembly graphs as well as a Python package interface to load and analyse assembly graphs from popular assemblers. *agtools* exposes a range of functionalities, including statistics generation, segment renaming, graph concatenation, segment filtering, component extraction and format conversion between multiple assembly graph formats, providing a unified solution for analysing assembly graphs. *agtools* can be easily integrated into bioinformatic tools and workflows that require assembly graph-based analyses.

*agtools* is implemented using a modular architecture, allowing individual components to be reused. It can be easily extended to add more functionality, enabling the addition of new graph operations and support for new formats. As future work, we plan to add support for more assemblers that generate assembly graphs and integrate more features such as graph-based sequence querying, which will broaden the utility of *agtools* in other genomic and metagenomic applications.

## Acknowledgements

This research was undertaken with the resources and services from the DeepThought HPC at Flinders University (Flinders University 2021), the Phoenix HPC at University of Adelaide, the National Computational Infrastructure (computing support was provided by an NCI Adapter allocation to V.M.) and the Pawsey Supercomputing Research Centre (computing support was provided by an NCMAS allocation to R.A.E.), which is supported by the Australian Government. We would like to thank Fabien Voisin and Sarah Beecroft for their assistance on Phoenix and Pawsey respectively. We acknowledge the use of ChatGPT (OpenAI) to assist in improving clarity and readability of the manuscript, code development and debugging, and the preparation of documentation. The authors reviewed all content generated with ChatGPT to confirm its accuracy and integrity.

## Author contributions

V.M. conceived the project, developed the software, performed all analyses and wrote the paper. V.M. and G.B. preprocessed the datasets. G.B., R.R.W., S.R.G. and B.P. tested the software. R.A.E. supervised the project. All the authors reviewed the manuscript and provided detailed feedback.

## Data and code availability

All the datasets containing raw sequencing data used for this work are publicly available from their respective studies on NCBI. The Tara Oceans datasets were downloaded from BioProject number PRJEB4419, the human gut metagenomes from BioProject number PRJNA820119 and Microflora Danica datasets from BioProject number PRJEB58634. The assembly outputs are available on Zenodo at https://zenodo.org/records/17075231 (short-read assemblies) and https://zenodo.org/records/17096649 (long-read assemblies).

The code of *agtools* is freely available on GitHub under the MIT license and can be found at https://github.com/Vini2/agtools. All tests in this study were performed using *agtools* v.1.0.0.

