## Supplementary Material for "*agtools*: a software framework to manipulate assembly graphs"


Vijini Mallawaarachchi[1*], George Bouras[2,3], Ryan R. Wick[4,5], Susanna R. Grigson[1], Bhavya Papudeshi[1], Robert A. Edwards[1]

[1] Flinders Accelerator for Microbiome Exploration, College of Science and Engineering, Flinders University, Bedford Park, Adelaide, SA 5042, Australia

[2] Adelaide Medical School, Faculty of Health and Medical Sciences, The University of Adelaide, Adelaide, SA 5005, Australia.

[3] The Department of Surgery – Otolaryngology Head and Neck Surgery, Central Adelaide Local Health Network, Adelaide, SA 5005, Australia.

[4] Department of Microbiology and Immunology, The University of Melbourne at the Peter Doherty Institute for Infection and Immunity, Melbourne, VIC 3000, Australia

[5] Centre for Pathogen Genomics, The University of Melbourne, Parkville, VIC 3010, Australia

* Corresponding author

### Feature comparison of existing tools

Table S1. Feature comparison of existing tools

| **Tool** | **Formats supported** | **Supports graph format* conversion?** | **API or CLI available?** | **Can parse assembler-specific contig graphs?** | **Link** |
|---|---|---|---|---|---|
| gfatools | GFA | Yes | CLI | No | https://github.com/lh3/gfatools |
| PyGFA | GFA | No | API | No | https://github.com/pmelsted/pyGFA |
| GFA for Ruby | GFA | No | API | No | https://github.com/lmrodriguezr/gfa |
| RGFA (Gonnella and Kurtz 2016) | GFA | No | API | No | https://github.com/ggonnella/RGFA |
| GFAKluge (Dawson and Durbin 2019) | GFA | No | API and CLI | No | https://github.com/edawson/gfakluge |
| GfaPy (Gonnella and Kurtz 2017) | GFA | No | API | No | https://github.com/ggonnella/gfapy |
| Pyfastg | FASTG | No | API | No | https://github.com/fedarko/pyfastg |

* Converting other assembly graph formats such as FASTG and ASQG to GFA

### Runs used for performance analysis

Table S2. Datasets used for profiling *agtools*

| **Study** | **Bioproject number** | **SRA accession numbers used in this work** |
|---|---|---|
| Tara Oceans (de Vargas et al. 2015) | PRJEB4419 | ERR2750826, ERR2750828, ERR2752143, ERR2752144, ERR2752145, ERR2752146, ERR2752147, ERR2752149, ERR2752150, ERR2752151, ERR2752153, ERR2752154, ERR2752160, ERR2752162, ERR2752163, ERR321018, ERR594355, ERR594360, ERR594361, ERR594362, ERR594370, ERR594371, ERR594372, ERR594375, ERR599357, ERR599362, ERR599370, ERR599383 |
| Human gut (Chen et al. 2022) | PRJNA820119 | SRR18490951, SRR18490961, SRR18491036, SRR18491148, SRR18491176, SRR18491204, SRR18491300, SRR18491309, SRR18491312, SRR18491319 |
| Microflora Danica (Sereika et al. 2025) | PRJEB58634 | ERR12040030, ERR11593880, ERR11561019, ERR11523645, ERR10750395 |

### Running time and memory usage of `rename`

Table S3. Running time and memory usage of `rename` subcommand

| **Dataset** | **Size of GFA file** | **Wall clock time** | **Peak memory usage** |
|---|---:|---:|---:|
| ERR2752151 | 2.6GB | 55.59s | 2.668GB |
| ERR2752163 | 2.5GB | 43.62s | 1.690GB |
| ERR2752162 | 2.2GB | 38.07s | 1.540GB |
| ERR2752160 | 2.0GB | 37.97s | 1.544GB |
| ERR2752150 | 1.7GB | 29.73s | 1.386GB |
| ERR2752154 | 1.5GB | 28.54s | 1.386GB |
| ERR2752145 | 1.4GB | 24.30s | 955.924MB |
| ERR2752149 | 1.4GB | 24.81s | 943.316MB |
| ERR2752152 | 1.1GB | 19.12s | 802.820MB |
| ERR2752153 | 1.2GB | 16.14s | 747.304MB |

### Running time and memory usage of `concat`

Table S4. Running time and memory usage of `concat` subcommand

| Number of GFA files concatenated | Total size of GFA files | Wall clock time | Peak memory usage |
|---|---:|---:|---:|
| 2 | 5.8GB | 1m 9.80s | 3.258GB |
| 3 | 8.2GB | 1m 36.02s | 3.404GB |
| 4 | 9.3GB | 2m 06.36s | 4.022GB |
| 5 | 10.5GB | 2m 20.46s | 6.413GB |
| 6 | 12.4GB | 2m 42.47s | 6.413GB |
| 7 | 14.1GB | 2m 55.50s | 6.414GB |
| 8 | 15.3GB | 4m 02.14s | 6.490GB |
| 9 | 16.3GB | 3m 17.66s | 6.731GB |
| 10 | 17.2GB | 3m 25.66s | 6.887GB |
| 11 | 18.0GB | 3m 34.74s | 7.266GB |

### Running time and memory usage of `clean`

Table S5. Running time and memory usage of `clean` subcommand

| Dataset | Size of GFA file | Size of FASTA file | Wall clock time | Peak memory |
|---|---:|---:|---:|---:|
| ERR12040030 | 940MB | 25GB | 5m 7.63s | 1.047GB |
| ERR11593880 | 782MB | 22GB | 4m 23.14s | 743.200MB |
| ERR11523645 | 1.004GB | 12GB | 2m 41.08s | 603.716MB |
| ERR11561019 | 286MB | 6.5GB | 1m 28.40s | 375.708MB |
| ERR10750395 | 47MB | 416MB | 13.56s | 143.412MB |

### Running time and memory usage of `fastg2gfa`

Table S6. Running time and memory usage of `fastg2gfa` subcommand

| **Dataset** | **Size of FASTG file** | **Wall-clock time** | **Peak memory usage** |
|---|---:|---:|---:|
| ERR2752163 | 3.4GB | 25.35s | 6.628GB |
| ERR2752147 | 3.1GB | 29.06s | 5.981GB |
| ERR2752151 | 2.8GB | 20.52s | 5.518GB |
| ERR2752146 | 2.5GB | 23.87s | 4.954GB |
| ERR2752150 | 2.3GB | 14.46s | 4.298GB |
| ERR2752145 | 2.0GB | 17.83s | 3.823GB |
| ERR2752149 | 1.9GB | 13.15s | 3.693GB |
| ERR2752153 | 1.6GB | 11.53s | 3.000GB |
| ERR2752144 | 1.5GB | 15.03s | 2.966GB |
| ERR2752143 | 1.2GB | 11.37s | 2.338GB |

### cProfile profiling results

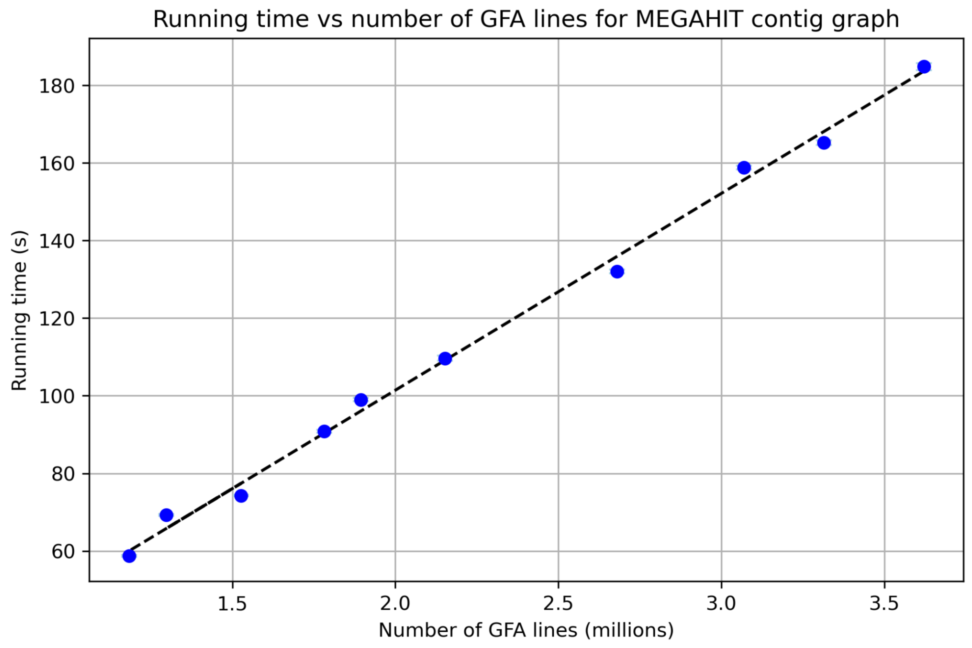


**Fig. S1** Running time to load contig-level assembly graphs from MEGAHIT assembler.

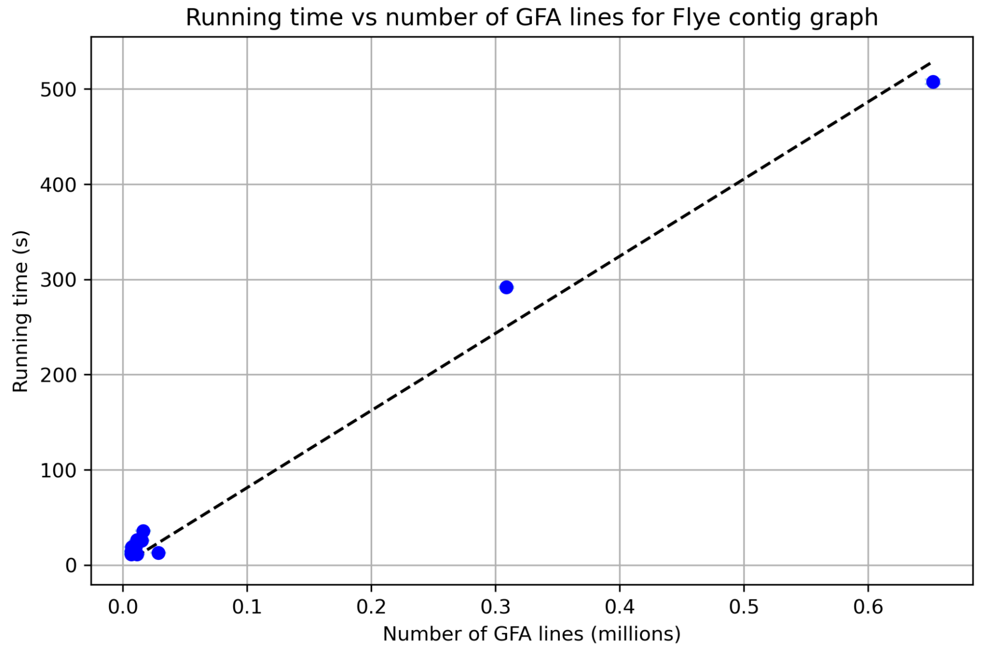


**Fig. S2** Running time to load contig-level assembly graphs from Flye assembler.

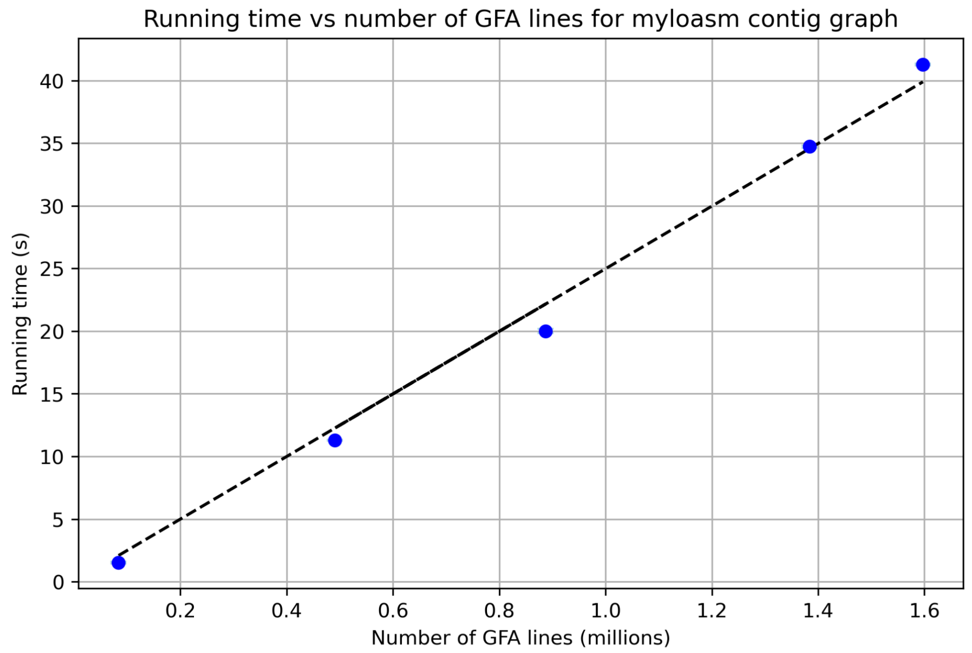


**Fig. S3** Running time to load contig-level assembly graphs from myloasm assembler

### Assemblers used

1. metaSPAdes (Nurk et al. 2017) from SPAdes (Bankevich et al. 2012)
2. MEGAHIT (Li et al. 2015)
3. metaFlye (Kolmogorov et al. 2020) from Flye (Kolmogorov et al. 2019)
4. myloasm (Shaw, Marin, and Li 2025)

Gonnella, Giorgio, and Stefan Kurtz. 2016. “RGFA: Powerful and Convenient Handling of Assembly Graphs.” *PeerJ* 4 (November):e2681.

———. 2017. “GfaPy: A Flexible and Extensible Software Library for Handling Sequence Graphs in Python.” *Bioinformatics (Oxford, England)* 33 (19): 3094–95.

Kolmogorov, Mikhail, Derek M. Bickhart, Bahar Behsaz, Alexey Gurevich, Mikhail Rayko, Sung Bong Shin, Kristen Kuhn, et al. 2020. “metaFlye: Scalable Long-Read Metagenome Assembly Using Repeat Graphs.” *Nature Methods* 17 (11): 1103–10.

Kolmogorov, Mikhail, Jeffrey Yuan, Yu Lin, and Pavel A. Pevzner. 2019. “Assembly of Long, Error-Prone Reads Using Repeat Graphs.” *Nature Biotechnology* 37 (5): 540–46.

Li, Dinghua, Chi-Man Liu, Ruibang Luo, Kunihiko Sadakane, and Tak-Wah Lam. 2015. “MEGAHIT: An Ultra-Fast Single-Node Solution for Large and Complex Metagenomics Assembly via Succinct de Bruijn Graph.” *Bioinformatics (Oxford, England)* 31 (10): 1674–76.

Nurk, Sergey, Dmitry Meleshko, Anton Korobeynikov, and Pavel A. Pevzner. 2017. “metaSPAdes: A New Versatile Metagenomic Assembler.” *Genome Research* 27 (5): 824–34.

Sereika, Mantas, Aaron James Mussig, Chenjing Jiang, Kalinka Sand Knudsen, Thomas Bygh Nymann Jensen, Francesca Petriglieri, Yu Yang, et al. 2025. “Genome-Resolved Long-Read Sequencing Expands Known Microbial Diversity across Terrestrial Habitats.” *Nature Microbiology* 10 (8): 2018–30.

Shaw, Jim, Maximillian G. Marin, and Heng Li. 2025. “High-Resolution Metagenome Assembly for Modern Long Reads with Myloasm.” *bioRxiv*. https://doi.org/10.1101/2025.09.05.674543.

Vargas, Colomban de, Stéphane Audic, Nicolas Henry, Johan Decelle, Frédéric Mahé, Ramiro Logares, Enrique Lara, et al. 2015. “Ocean Plankton. Eukaryotic Plankton Diversity in the Sunlit Ocean.” *Science (New York, N.Y.)* 348 (6237): 1261605.
